# A non-structural pure enzyme protein forms a LCST type of stimuli-responsive and reversible hydrogel with novel structure and catalytic activity

**DOI:** 10.1101/2021.02.07.430034

**Authors:** J. Nie, X. Zhang, Y. Liu, M.A. Schroer, W. Wang, J. Ren, D.I. Svergun, A.-P. Zeng

**Affiliations:** Beijing Advanced Innovation Center for Soft Matter Science and Engineering, Beijing University of Chemical Technology, North Third Ring Road 15, Chaoyang District, 100029, Beijing, China; Institute of Bioprocess and Biosystems Engineering, Hamburg University of Technology Denickestrasse 15, D-21073 Hamburg, Germany; European Molecular Biology Laboratory (EMBL), Hamburg Outstation c/o DESY, Notkestrasse 85, 22607 Hamburg, Germany; State Key Laboratory for Biology of Plant Diseases and Insect Pests/Key Laboratory of Control of Biological Hazard Factors (Plant Origin) for Agri-product Quality and Safety, Ministry of Agriculture, Institute of Plant Protection, Chinese Academy of Agricultural Sciences, Beijing 100081, China

**Author notes:** Corresponding author: Prof. An-Ping Zeng. Authors contributed equally (J. Nie, X. Zhang).

**Keywords:** Protein, Enzyme, Hydrogel, Lipoate-protein ligase, Small angle X-ray scattering, Phase transition, Crosslinking network

## Abstract

Hydrogels have a wide range of applications such as in biomedicine, cosmetics and soft electronics. Compared to polymer hydrogels based on covalent bonding, protein hydrogels offer distinct advantages owing to their biocompatibility and better access to molecular engineering. However, pure and natural protein hydrogels have been seldom reported except for structural proteins like collagen and silk fibrin. Here, we report the unusual ability and mechanism of a unique natural enzyme, lipoate-protein ligase A (LplA) of *E. coli* to self-assemble into a stimuli-responsive and reversible hydrogel of the low critical solution temperature (LCST) type. This is the first globular and catalytic protein found to form a hydrogel in response to temperature, pH and the presence of ions. Protein structure based analysis reveals the key residues responsible for the gel formation and mutational studies confirms the essential roles of hydrogen bonding between the C-terminal domains and electrostatic interactions in the N-terminal domains. Characterization of phase transitions of wild type LplA and its mutants using small angle X-ray scattering (SAXS) yields details of the gelation process from initial dimer formation over a pre-gel-state to full network development. Further electron microscopic analyses and modeling of SAXS data suggest an unusual interlinked ladder-like structure of the macroscopic crosslinking network with dimers as ladder steps. The unique features of this first reported protein hydrogel may open up hitherto inaccessible applications, especially those taking advantage of the inherent catalytic activity of LplA.

## Introduction

As macromolecular soft matter, hydrogels have received great interests for applications in biomedicine like tissue engineering (Murphy and Atala 2014; Todhunter et al. 2015), controlled drug release (Gupta et al. 2017; Li and Mooney, 2017; Liang et al. 2019), and wound healing (Moore et al. 2018; Lopez-Silva et al. 2020). New applications are emerging in many other areas such as soft electronics (Lin et al. 2016), biosensors and actuators (Wang et al. 2010; MacConaghy et al. 2014), agriculture (Lu et al. 2016; Xiang et al. 2018) and food industry (McClements 2015; Cao and Mezzenga 2020). Hydrogels are special aqueous dispersion systems comprised of three-dimensional (3D) networks of polymers, which are hydrophilic but insoluble as a result of the presence of cross-linking interactions among the constituents (Zhang and Khademhosseini 2017). According to the source of the constituents, they can be classified into synthetic, natural and hybrid hydrogels. Another classification is according to the type of cross-linking interactions, which can be either chemical or physical. Hydrogels cross-linked by covalent bonds exhibit outstanding mechanical properties, but some of the cross-linking agents are toxic to cells (Jonker et al. 2012), which limits the potential applications as medical materials. Further, covalent bonding of polymers is normally very stable, and the transition from sol to gel is therefore generally irreversible (Zhang and Khademhosseini 2017; Fang et al. 2013). In comparison, physically cross-linked hydrogels are assembled through non-covalent bonds, such as hydrogen bonding, electrostatic interaction and hydrophobic interaction, which lead to weaker but reversible, and less-toxic gels (Chassenieux and Tsitsilianis 2016; Shmidov et al. 2019).

Of the various hydrogels reported, protein-based hydrogels have several distinct advantages owing to their well-defined sequences, biocompatibility, stimuli-response and adjustable properties (Jonker et al. 2012; Banta et al. 2010; Kopecek and Yang 2012; Wang et al. 2013). The past two decades have seen increasingly rapid development of protein-based functional hydrogels used in tissue engineering (Wang et al. 2013; Roberts et al. 2018; Yamada et al. 2019), drug delivery (Saxena and Nanjan 2015; Hill et al. 2019; Yamada et al. 2019), and 3D cell culture (Gelain et al. 2006; Wu et al. 2011; Jeon et al. 2011; Yang et al. 2019; Mizuguchi et al. 2020). They are usually physically cross-linked hydrogels, and interactions between peptides or protein domains are the main driving force for the self-assembly (Jonker et al. 2012; Banta et al. 2010; Kopecek and Yang 2012). The representative protein building blocks include coiled-coil (Petka 1998; Wang et al. 1999; Jing et al. 2008; Olsen et al. 2018; Hill et al. 2019), β-hairpin peptide (Collier et al. 2001; Schneider et al. 2002; Pochan et al. 2003; Kretsinger et al. 2005; Wu et al. 2011; Yamada et al. 2019;), silk fibrin (Kim et al. 2004; Buitrago et al. 2018), SpyCatcher-SpyTag (Lyu et al. 2017; Wang et al. 2017; Yang et al. 2019; Wang et al. 2019), WW domain-polyproline containing peptide (Maciasa et al.2002; Foo et al. 2009), and TIP-1:Kir complex (Guan et al. 2013; Xu et al. 2017; Wu et al. 2018). Although nature may provide a large number of functional protein building blocks capable of forming hydrogels, only a small part of them has been discovered and explored. To our best knowledge, no other proteins or peptides with low critical solution temperature (LCST) have been reported up to now, except for the structural protein elastin, which has been extensively studied (Urry 1992; Wright and Conticello 2002; Lao et al. 2008; Saxena and Nanjan 2015; Tarakanova et al. 2017; Roberts et al. 2018; Mizuguchi et al. 2020). In the field of biomedicine, there is a great interest on thermoresponsive hydrogels or modules for applications under near physiological conditions.

We report here the unusual ability and mechanism of a unique natural enzyme, lipoate-protein ligase A (LplA) of *Escherichia coli* to self-assemble into a reversible hydrogel. LplA is responsible for the lipoylation of many important proteins (enzymes) such as the E2 subunit of pyruvate dehydrogenase and 2-ketoglutarate dehydrogenase complexes, and the H-protein of the glycine cleavage system (GCS). GCS is involved in the one-carbon metabolism of all types of cells (Kikuchi et al. 2008; Zhang et al. 2019; Zhang et al. 2020a; Zhang et al. 2020b; Xu et al. 2020), among others in photorespiration of plants (Timm et al. 2018; Lopez-Calcagno et al. 2019) and disease development including aging, obesity and cancers (Amelio et al. 2014; Zhang et al. 2012; Gao et al. 2019). We proved that it still maintains *in vitro* catalytic activity in the hydrogel state. It is the first natural protein and also the first globular protein which is found to form a hydrogel with tuneable LCST in its pure form. LplA-based hydrogels offer the decisive advantages of being composed of a catalytic enzyme compared to all other known hydrogels and may therefore open up hitherto inaccessible applications (Tang et al. 2020).

## Results and Discussion

### Catalytic hydrogel with LCST behavior formed by pure LplA

We discovered that His-tag purified LplA from *E. coli* exhibits a reversible LCST behaviour in Tris-HCl buffer (50 mM, pH 7.5), cycling between a clear solution and a turbid hydrogel (**Fig.1a**). A stable hydrogel can be observed at room temperature (*T* = 20°C) when the protein concentration is above 40 mg/mL. The presence of His-tag has no effect on the phase behaviour, as shown by examining LplA protein with His-tag cleaved off under the same experimental conditions (**Fig. 1a)**. The opacity of the gel formed by *E. coli* LplA suggests that protein aggregation (or microscopic phase separation) occurs during gelation. The formation of reversible aggregates is generally considered to be caused by self-assembly of proteins (Mahler et al. 2009), induced by non-covalent interactions between the molecules.

**Figure 1.**
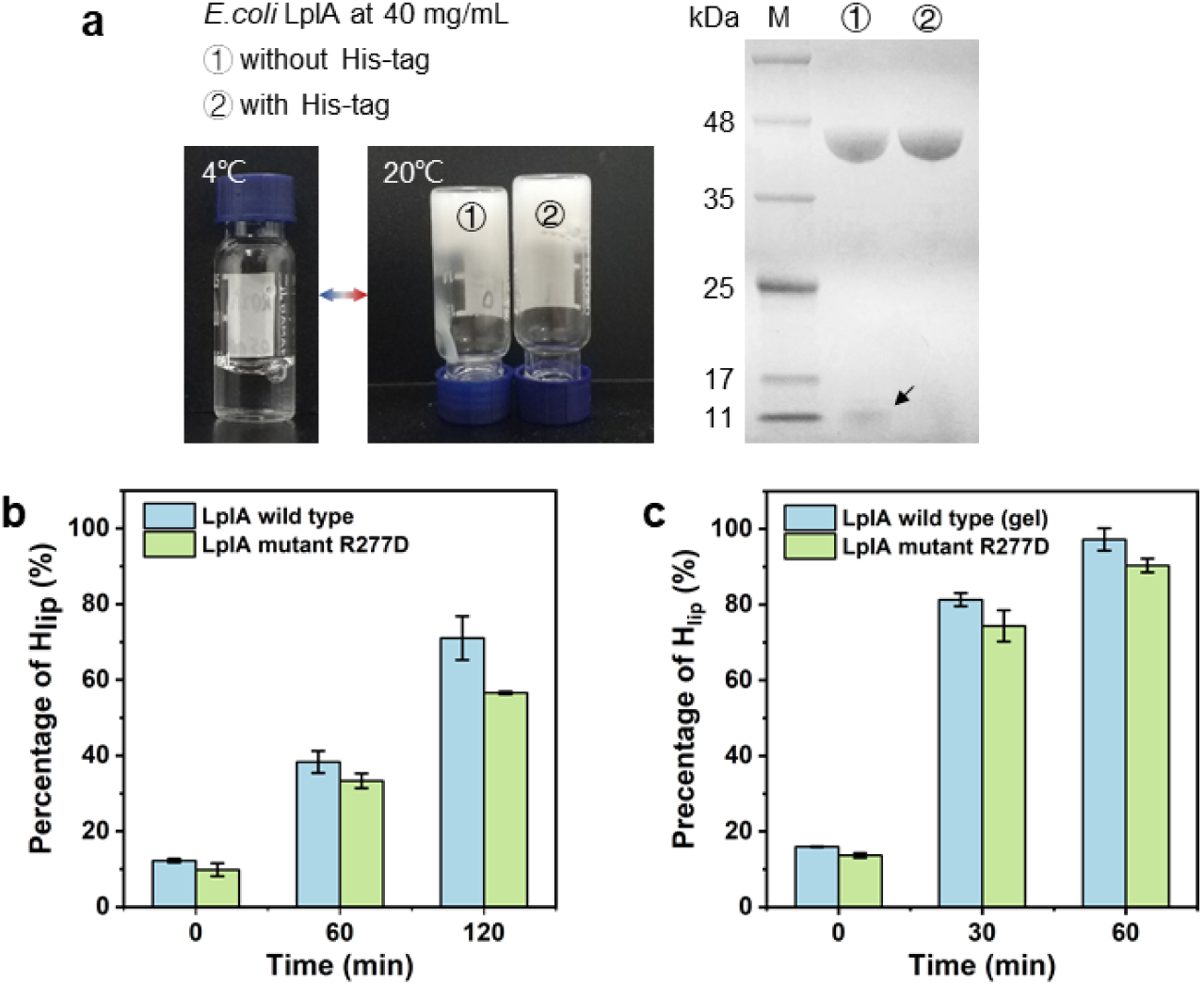
Catalytic hydrogel of *E. coli* LplA with LCST behaviour. **a**. Thermoresponsive cycling of LplA between a solution phase and a gel phase. At a concentration of 40 mg/mL, LplA solution with or without His-tag can undergo phase separation and form hydrogel by increasing the temperature. 12% SDS-PAGE showed cleaved His-Tag in group① at 11 kDa (arrow). **b**. Catalytic activities of LplA wild type and mutant R277D in a solution for the lipoylation of H-protein of the glycine cleavage systems. The concentration of LplA is 0.8 mg/mL. **c**. Catalytic activities of LplA wild type in a gel state and mutant R277D in solution. The concentration of LplA is 40 mg/mL.

In order to examine whether LplA still has catalytic activity in the gel form, we determined the lipoylation of H-protein at low and high LplA concentrations. As a comparison, a mutant R277D, which cannot form a gel at 30°C (see below), was also tested. As shown in **Fig. 2b**, the mutation of R277 did not significantly affect the catalytic activity. When at high protein concentrations (4o mg/ml), the LplA wild type (in a gel state) and the mutant R277D (in a solution state) show nearly the same lipoylation rate of H-protein **(Fig. 2c)**. These results confirm that LplA retains its catalytic activity in the gel state.

**Figure 2.**
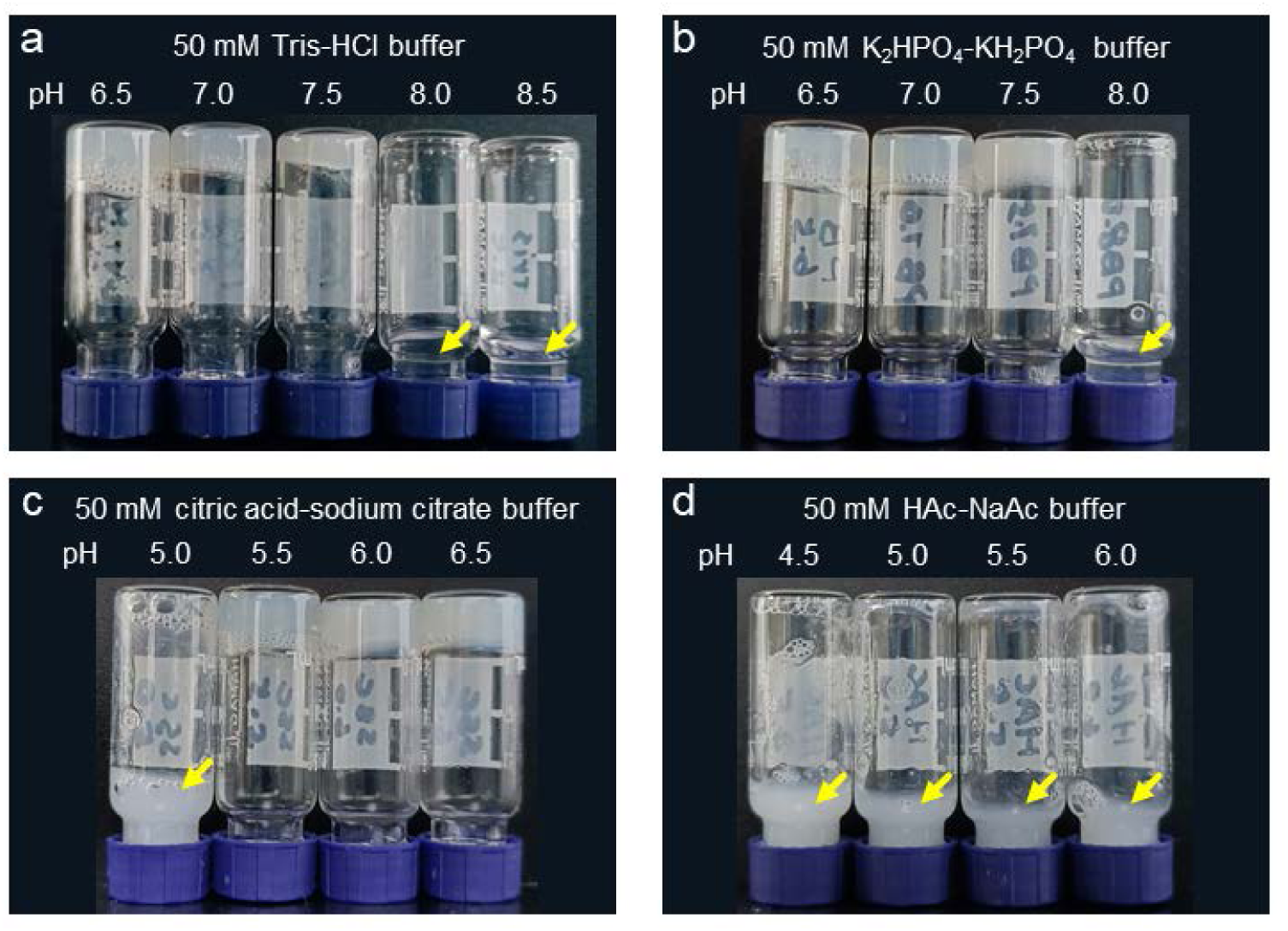
Formation of LplA hydrogel at room temperature (20°C) in different buffer solutions with different pH values (protein concentration = 40 mg/mL). Samples failing in gel formation are marked by arrows.

Further examinations of LplA in other typical buffers reveal a sharp pH response and a dependence on buffer ions (**Fig.2**). In a Tris-HCl buffer or K_2_HPO_4_-KH_2_PO_4_ buffer, LplA forms a reversible hydrogel in the pH range of 6.5∼7.5 (**Fig. 2a, 2b**), whereas at higher pH values (>8.0) LplA presents in a clear solution. In a citric acid-sodium citrate buffer, hydrogel can form in the lower pH range of 5.5∼6.5 (**Fig. 2c**); however, when the pH is below 5.0, LplA aggregates irreversibly, forming an emulsion instead of a gel. In an acetic acid-sodium acetate buffer, only a turbid LplA solution is formed (**Fig. 1d**). In the following experiments, unless otherwise specified, the solution environment was selected to be Tris-HCl buffer (pH 7.5).

### Dynamic rheological properties

Macroscopic mechanical properties of the LplA hydrogel were assessed via rheological analysis of the storage and loss modulus (G′ and G″, respectively). Strain sweep assessment at 30°C revealed a linear viscoelastic region below 0.1% strain (**Fig. 3a**), suggesting that the LplA hydrogel cannot withstand large shear strains. A strain of 0.1% was, therefore, used for frequency sweeps. In the tested frequency range, the sample is mainly elastic with the G′ values being nearly constant and exceeding the G″ values (**Fig. 3b**). Temperature sweep shows that the transition temperature (*T*_*t*_) is around 13°C, and the solution turnsd into a gel state above 20°C (**Fig. 3c**). To evaluate the ability of the hydrogel to reform after shear-thinning, we conducted a continuous ramp of shear rate experiment. Shear-thinning can be observed when the shear rate reaches 0.7 s^-1^. When the shear rate is restored to the minimum, the strength of the gel does not return to the original level (**Fig. 3d**). Interestingly, after the hydrogel re-experienced the cycle from low temperature to high temperature, its strength can be recovered completely.

**Figure 3.**
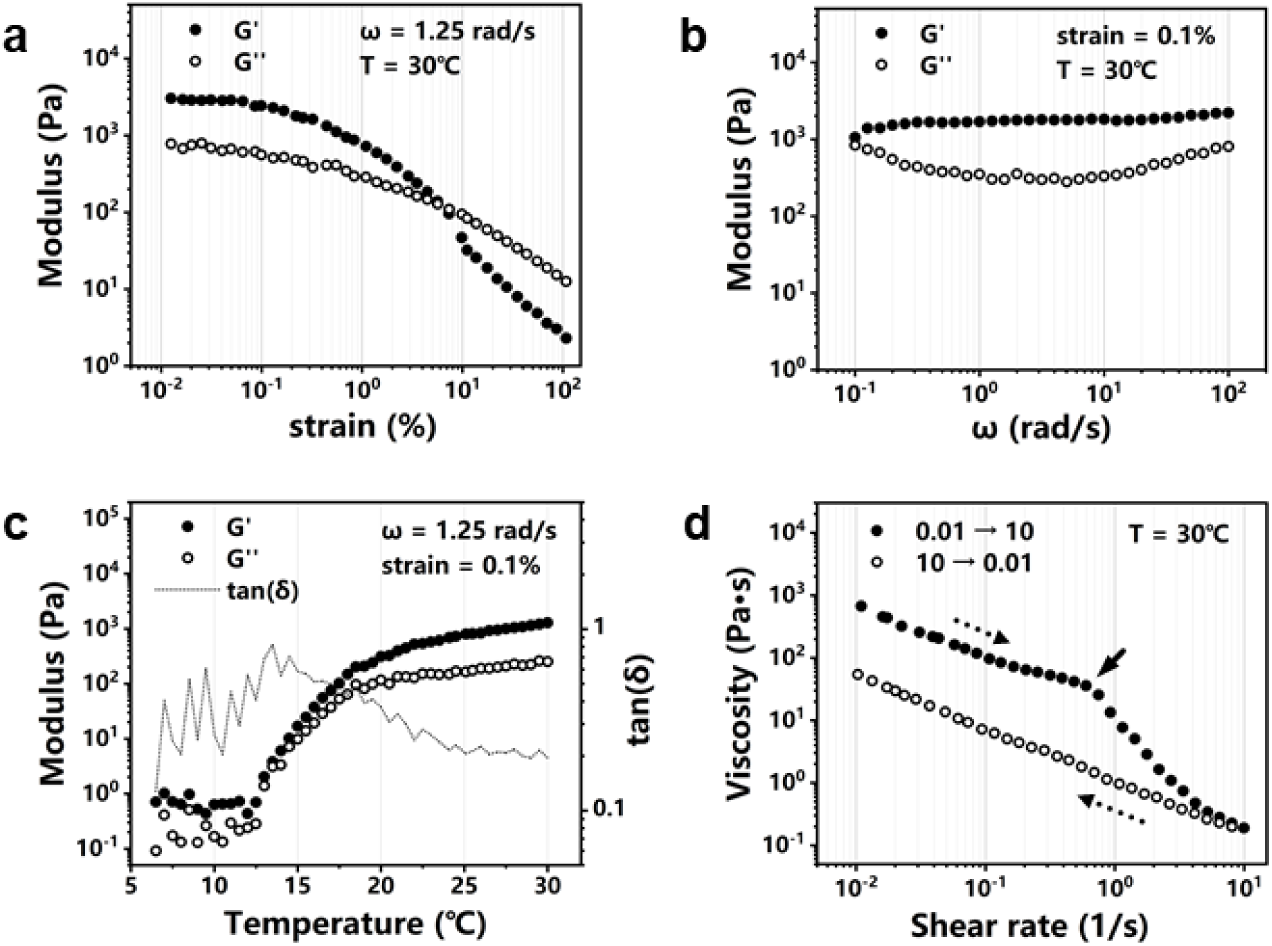
Rheological characterization of LplA hydrogel at a protein concentration of 40 mg/mL. **a**. Strain sweep at 30°C with a frequency constant of 1.25 rad/s. **b**. Angular frequency sweep at 30°C with a strain constant of 0.1%. **c**. Temperature sweep with a frequency constant of 1.25 rad/s and a strain of 0.1%. **d**. Continues ramp at 30°C°C with shear rate increased from 0.01 to 10 s^-1^ in 10 min and then dropped back to 0.01 s^-1^ in another 10 min.

### Hydrogel macroscopic crosslinking network

Aggregates formed by self-assembly of LplA molecules are also evident through transmission electron microscopy (TEM) measurement (**Fig 4a**). The images showed that the aggregates exhibit a coral-like porous structure with pore sizes of 5-10 nm. Obviously different from the physical entanglement of fiber assembly (Hill et al. 2019), the strength of the LplA hydrogel is mainly contributed by intermolecular interactions, which cannot be recovered quickly and completely after being sheared, as shown in the rheological test (**Fig. 3d**). This is probably because of the formation of LplA dimer as building block for the formation of gel via rather local interactions as revealed by our structure-based analysis and characterization of the gel formation using small angle X-ray scattering (SAXS) described below.

**Figure 4.**
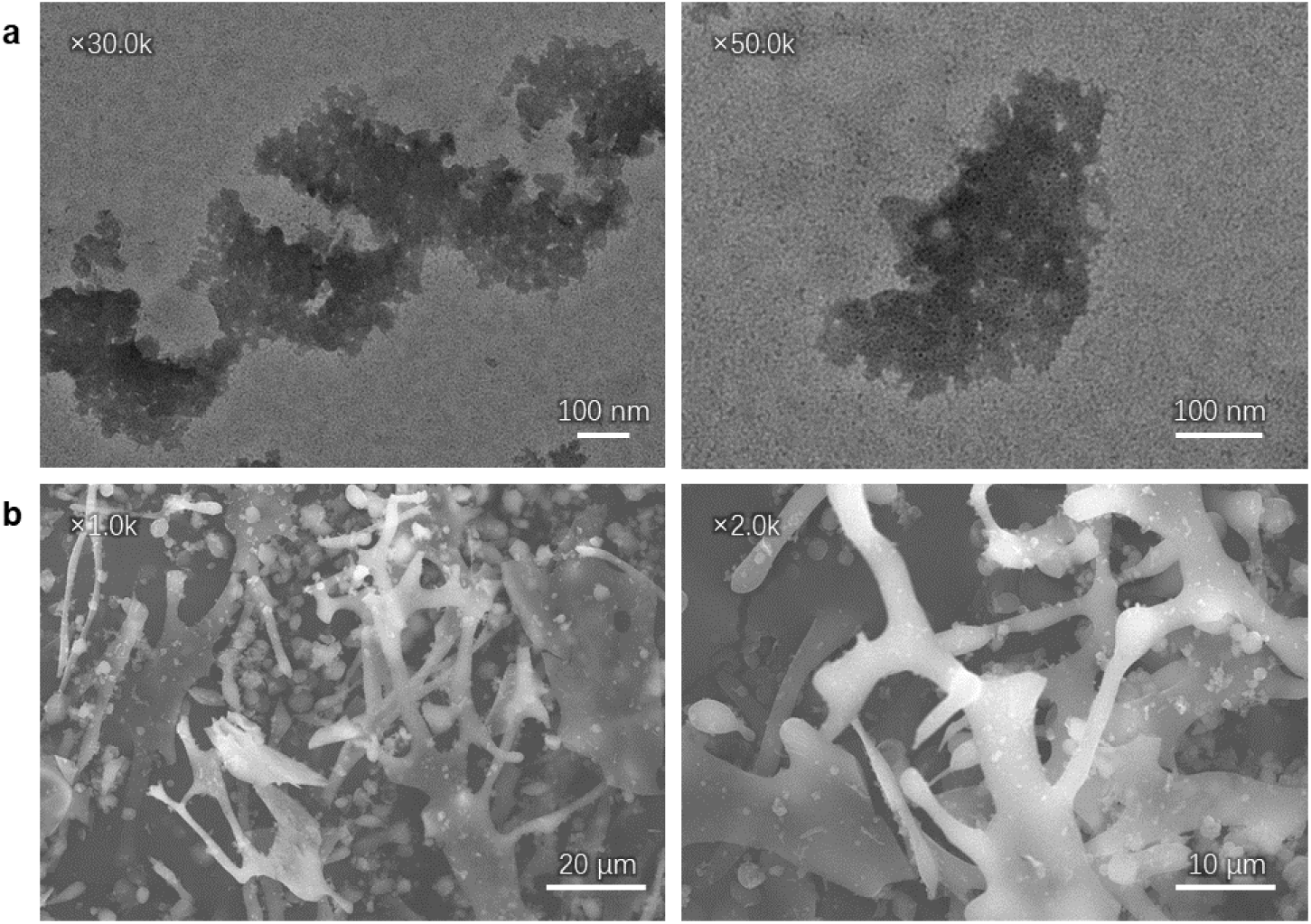
**a**. TEM images of *E. coli* LplA aggregate (magnification 30.0k and 50.0k respectively). **b**. SEM images of *E. coli* LplA hydrogel at a concentration of 40 mg/mL (magnification 1.0k and 2.0k respectively).

Scanning electron microscopy (SEM) was also employed to probe the microstructure of the LplA hydrogel samples. Although the gel network in the field of imaging appears fragmented due to its frangibility (**Fig 4b**), dendritic and lamellar network structures can still be observed. In addition, there are many globular protein particles in the background. Thus, we speculate that the microstructure of the gel network is formed by the fusion of phase-separated protein droplets.

### Structure-based analysis and mutants screening

Most of the reported structures of LplA homologues consist of a large N-terminal domain and a small C-terminal domain. However, unlike other homologues, *E. coli* LplA can undergo large-scale conformational changes between a bending conformation and a stretched conformation (**Fig. 5a**) (Fujiwara et al. 2005; Fujiwara et al. 2010). Such a conformational flexibility may be beneficial to the formation of a larger and more stable molecular interaction interface. In the crystal structure 3A7A, LplA exists as a dimer. At the surface of the C-terminal domain, there are dense polar residues, which can act as donors or acceptors for hydrogen bonding, and dipole-dipole interactions (**Fig. 5b**). Therefore, gelation of the individual single-point mutant of these polar residues at 20°C was tested (**Fig. 5d**). The transition temperature *T*_*t*_ is at around 14°C for S252A, D269A and Q279A, 16°C for H267A, and 24.5°C for R277A, respectively. Rheological experiments show that the gel strength of R277A is weaker than that of the wild type, while H267A forms unexpectedly stronger gel than the wild type (**Fig. 6a, 6b vs. Fig. 3c**). We therefore speculate that the interactions between the C-terminals of two LplA molecules play a key role in the protein aggregation process, and H267 and R277 have the most significant effects by forming hydrogen bonding.

**Figure 5.**
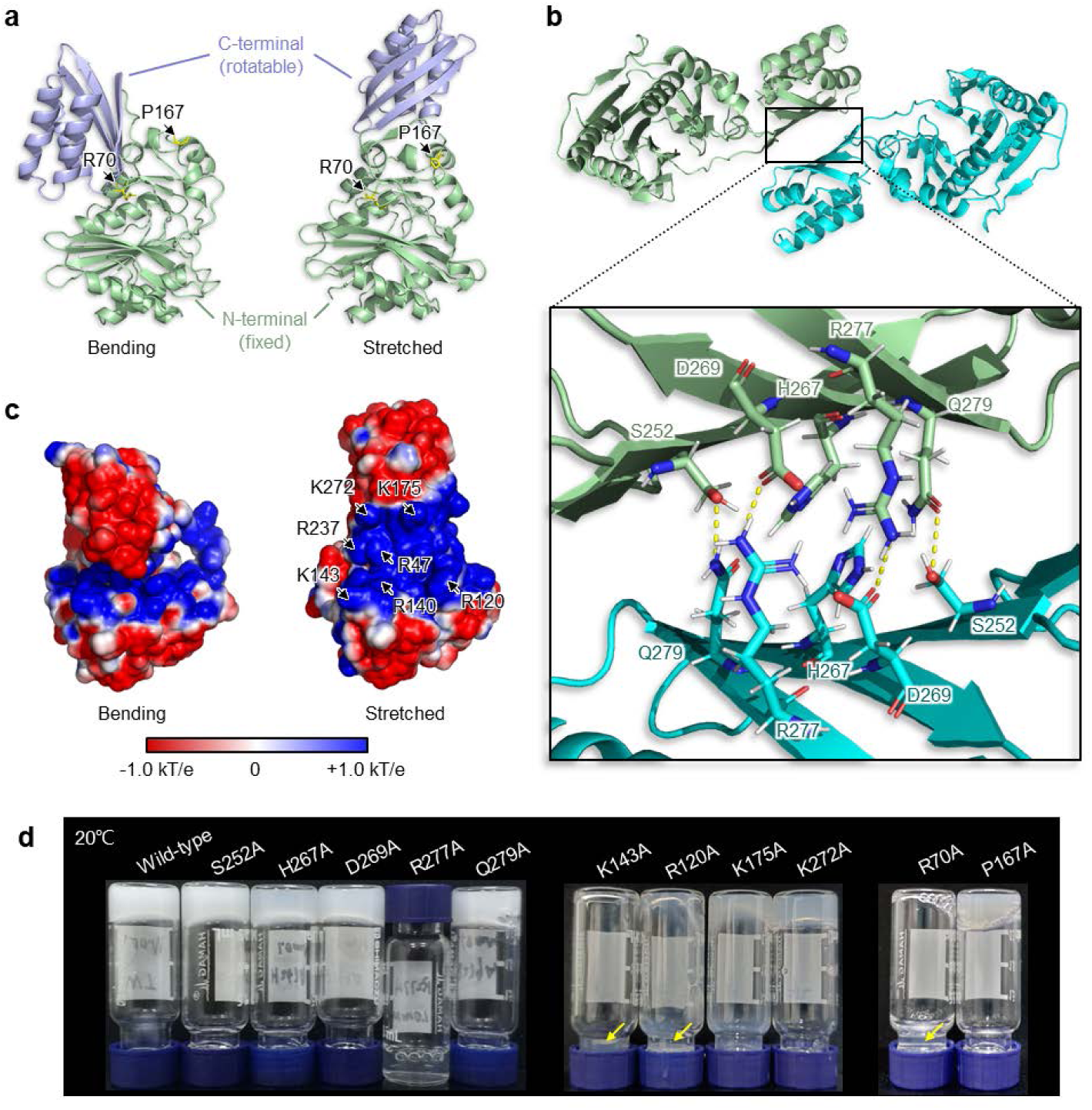
Structure-based analysis of mutants screening of *E. coli* LplA. **a**. Bending conformation (PDB IDs: 1×2G and 1×2H) and stretched conformation (PDB IDs: 3A7A and 3A7R) of *E. coli* LplA. **b**. Predicted molecular interactions between two C-terminal domains of *E. coli* LplA dimer (PDB ID: 3A7A). **c**. Projection of electrostatic potential isosurface on protein surface. The electrostatic potential is shown in the range of +1.0 kT/e (blue) and −1.0 kT/e (red). A large area with positive charge is exposed in the stretched conformation, which is related to the dense arginine and lysine. **d**. Gelation of *E. coli* LplA and its mutants at 20°C.

**Figure 6.**
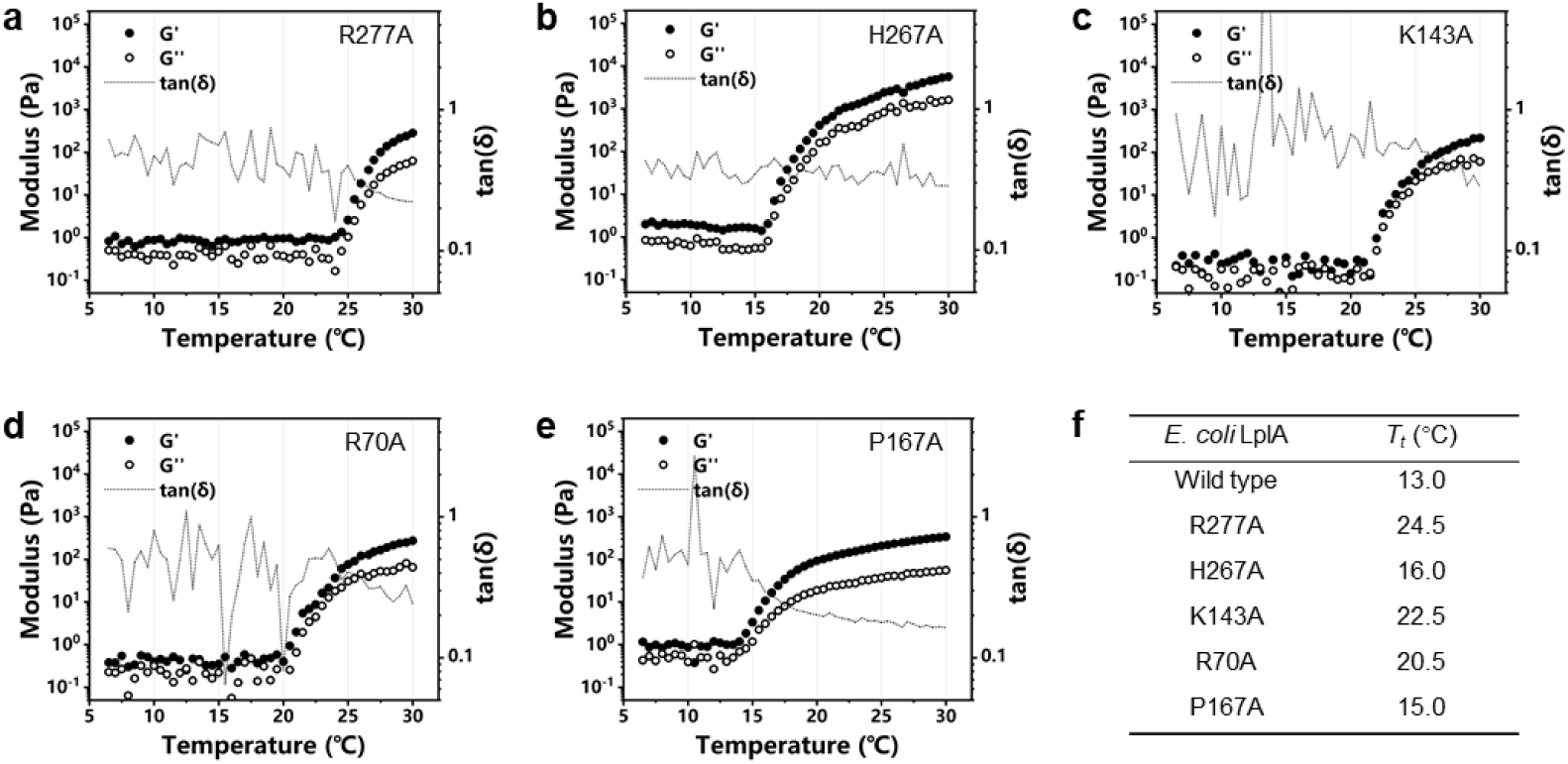
Temperature sweep of mutants **a**. R277A **b**. H267A **c**. K143A **d**. R70A **e**. P167A with frequency constant at 1.25 rad/s and strain at 0.1%. **f**. A summary of the transition temperature (*T*_*t*_) of wild type LplA and its mutants.

Another feature of the LplA molecule is the condensed presence of arginine and lysine in the H-protein binding domain in the N-terminal domain. The calculation of the electrostatic potential on the surface of LplA protein shows that this domain is completely covered by positive charges (**Fig. 5c**). In order to determine the influence of this domain on protein assembly, we selected four amino acid residues located on the boundary of the region for point mutations - R120A, K143A, K175A and R272A (**Fig. 5c**). The gelation of these mutants shows that K143 has the greatest impact on the stability of the gel (**Fig. 5d**). No gel formation of this mutant can be observed at 20°C. Its *T*_*t*_ is increased to 22.5°C (**Fig. 6f**).

R70 is located on the loop domain outside of the substrate pocket, which stabilizes the local protein conformation (**Fig. 5a**) (Fujiwara et al. 2005). P167 is located at the end of the flexible loop of LplA, which fixes the direction of the loop rotation (**Fig. 5a**). Their point mutants show weaker gel strength than the wild type (**Fig.5d**), and their transition temperatures also shift to higher values (**Fig. 6d, 6e and 6f**).

### Characterization of phase transition by small angle X-ray scattering (SAXS)

SAXS measurements were performed on wild type LplA and five of its mutants (R277A, H267A, K143A, R70A and P167A) under different concentrations. **Fig. 7a** depicts the SAXS curves *I*(*s*) of LplA at a concentration of *c* = 40 mg/mL during the heating cycle (scattering vector *s* = 4π/λ sin(Θ), with X-ray wavelength λ and scattering angle 2Θ). For temperatures *T* ≤ 12°C°C, the SAXS curves reveal the presence of monomeric LplA, as also indicated by the estimated molecular mass (**Tab. S1**), and unspecific, temperature-independent aggregates, as revealed by the intensity increase at smallest scattering angles and by comparing with the computed curve of the LplA monomer from the available crystal structure (dashed line; bending conformation). From 14°C onwards, the sample starts to change its structure. The forward scattering increases, revealing the formation of larger particles in the solution. Beginning from *T* = 20°C, distinct peaks evolve, heralding the formation of an ordered gel. The position of these correlation peaks, which reflect a low-range order within the gel, does not change significantly with temperature. These peaks can be subdivided into two sets. The first set, marked by dots, can be identified by broad Bragg reflections of a lamellar arrangement with a spacing of *d* ≈ 8.8 nm. The lamella structure is rather ordered and up to seven peaks can be distinguished. The second ordering consists of a single broad peak (diamond marker) at wave vector *s*_peak_ ≈ 1.0 nm^-1^, which corresponds to a distance of *d* = 2π/ *s*_peak_ ≈ 6.3 nm. At *T* = 30°C, these peaks are fully developed and do not change anymore, while the overall scattering intensity still increases. This indicates that the local order within the gel is fully established, while the gel continues to grow further in size.

**Figure 7.**
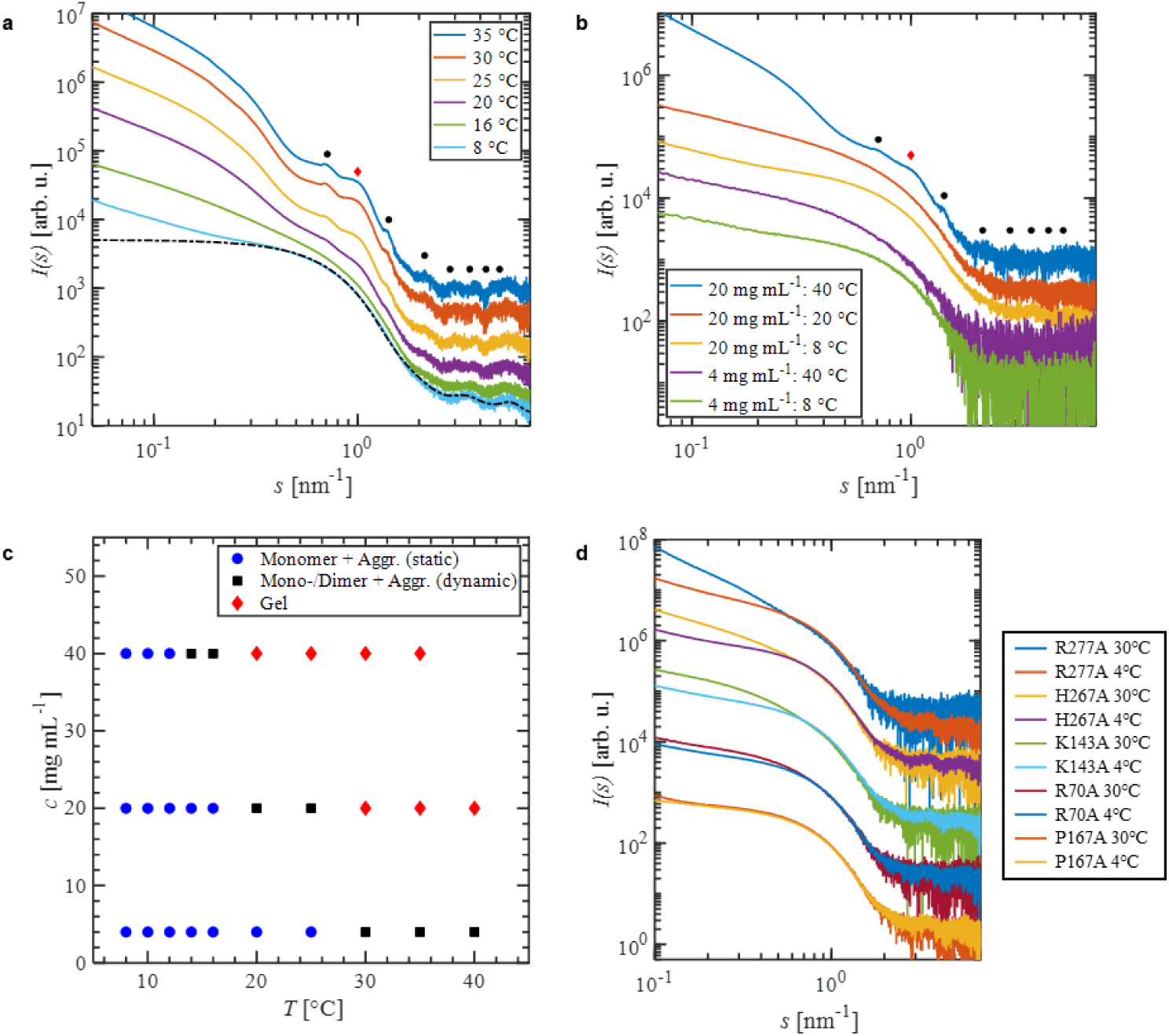
SAXS characterization of phase transition in the course of LplA hydrogel formation. **a**. SAXS profiles of LplA (c = 40 mg/mL) at different temperatures. At 25°C, correlation peaks appear, which are wholly developed at 30°C. The positions of the peaks are marked by dots (lamella order) and diamonds. The dashed line is the computed SAXS curve from the crystal structure of the bending conformation. The curves were displaced along the logarithmic axis for clarity. **b**. SAXS curves for lower LplA concentrations at the lowest and highest temperature measured. Symbols mark the position of correlation peaks. The same positions are present as for *c* = 40 mg/mL. Curves were shifted for clarity. **c**. Temperature-concentration phase diagram of LplA gel formation based on the SAXS measurements. **d**. SAXS profiles for five mutants at the lowest (*T* = 12°C) and highest (*T* = 30°C) temperature measured. Curves were shifted for clarity.

The SAXS data in the lowest temperature range can be reasonably well described by monomeric scattering with the presence of some unspecific aggregates. The temperature response can be divided into three regimes: monomer phase – aggregate formation – gel formation. Decreasing protein concentration results in a shift of the respective transition temperature. For *c* = 20 mg/mL, an ordered gel is present at *T* = 40°C, as can be seen from the Bragg peaks (**Fig. 7b**). While these peaks are not as pronounced as for the high concentration samples, the peak positions are the same, revealing the same spatial arrangement. For a concentration of *c* = 4 mg/mL, only the transition from the monomer to the aggregate is present in the temperature range measured. The concentration dependence suggests a diffusion-limited process, in which LplA monomers first interact with each other and then form higher order constructs. A tentative temperature – concentration phase diagram of the gelation process is given in **Fig. 7c**.

In order to understand the gel-formation process in more detail, the SAXS profiles of LplA at the different temperature regimes were modeled using the available high resolution models of the protein as building blocks (**Fig**.**8a**). For *T* ≤ 12°C, the temperature-independent data can be well described by the high-resolution structure of LplA in an stretched open conformation and allowing for flexibility of the C-terminal (**Fig. 8c)**; deviations at the smallest angles (*s* < 0.4 nm^-1^) stem from unspecific aggregates. Note that the crystal structures of the stretched conformation (PDB ID: 3A7R) as well as the bending conformation (PDB ID: 1×2G) of LplA cannot adequately describe the SAXS data (**Fig. 8b**). The SREFLEX analysis indicates that LplA is partially bent in the solution (**Fig.9a**). This partially bent conformer is thus utilized in the subsequent modeling. Starting from *T* = 14°C, larger species are observed, which can be well described by the formation of dimers for *s* > 0.4 nm^-1^ and larger unspecific aggregates.

**Figure 8.**
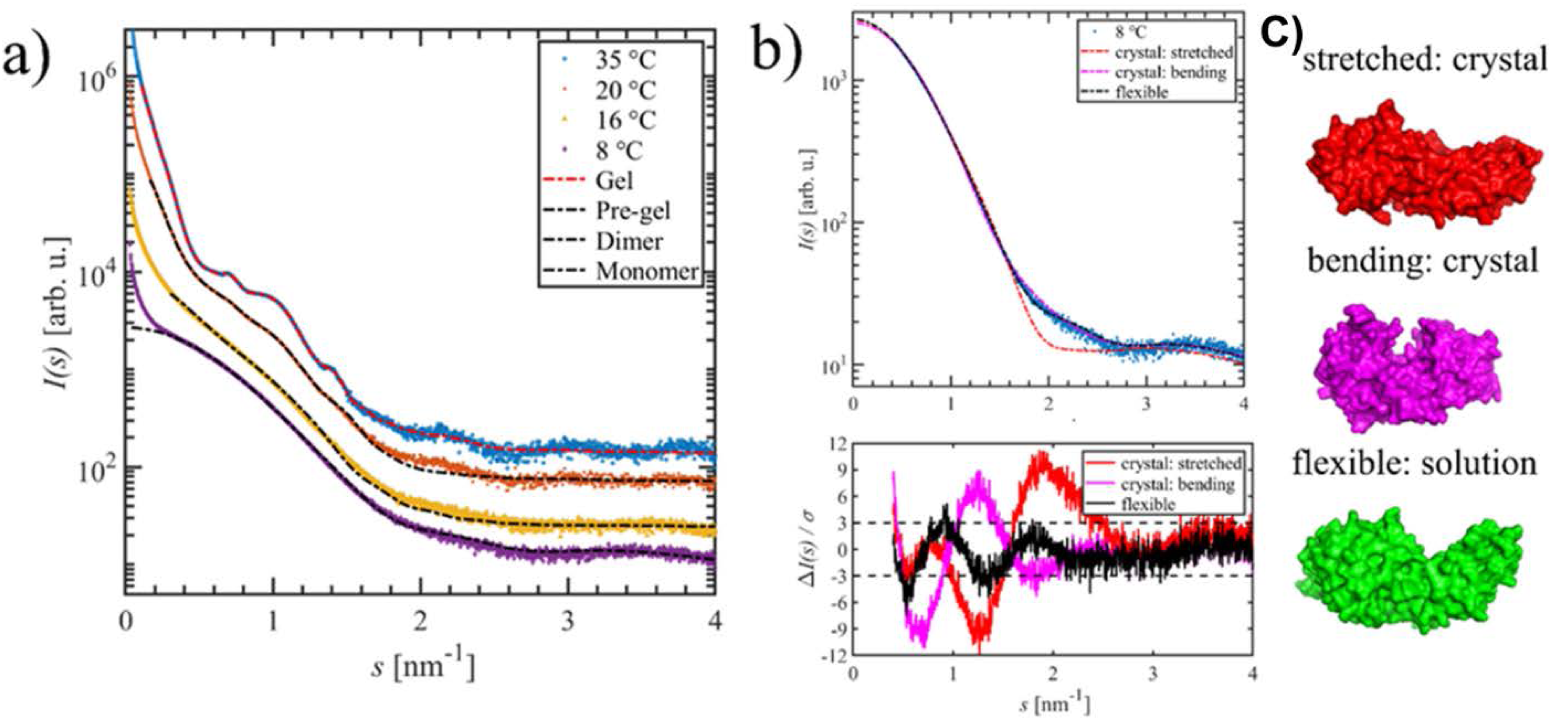
**a**. SAXS profiles of LplA at the different temperature regimes together with the fitting curves of the models: Monomer model of the high-resolution model including flexibility; dimer model using the flexible monomer and contacts at residues H267 between both units; pre-gel model using five of the dimer models and contact at residues K143 between the subunits; cluster model for the ordered gel, using 15 of the dimer models and K143 contacts. **b. Top:** Experimental SAXS curve of LplA and model curves for the stretched (PDB 3A7R) and bending (PDB 1×2G) crystal structure and when allowing for flexibility. **Bottom:** Residuals for the three fitting curves. Only the flexible model allows to describe the solution structure reasonably. **c**. Related models. The flexible solution state appears as an intermediate between the stretched and bending state present in the crystal.

**Figure 9.**
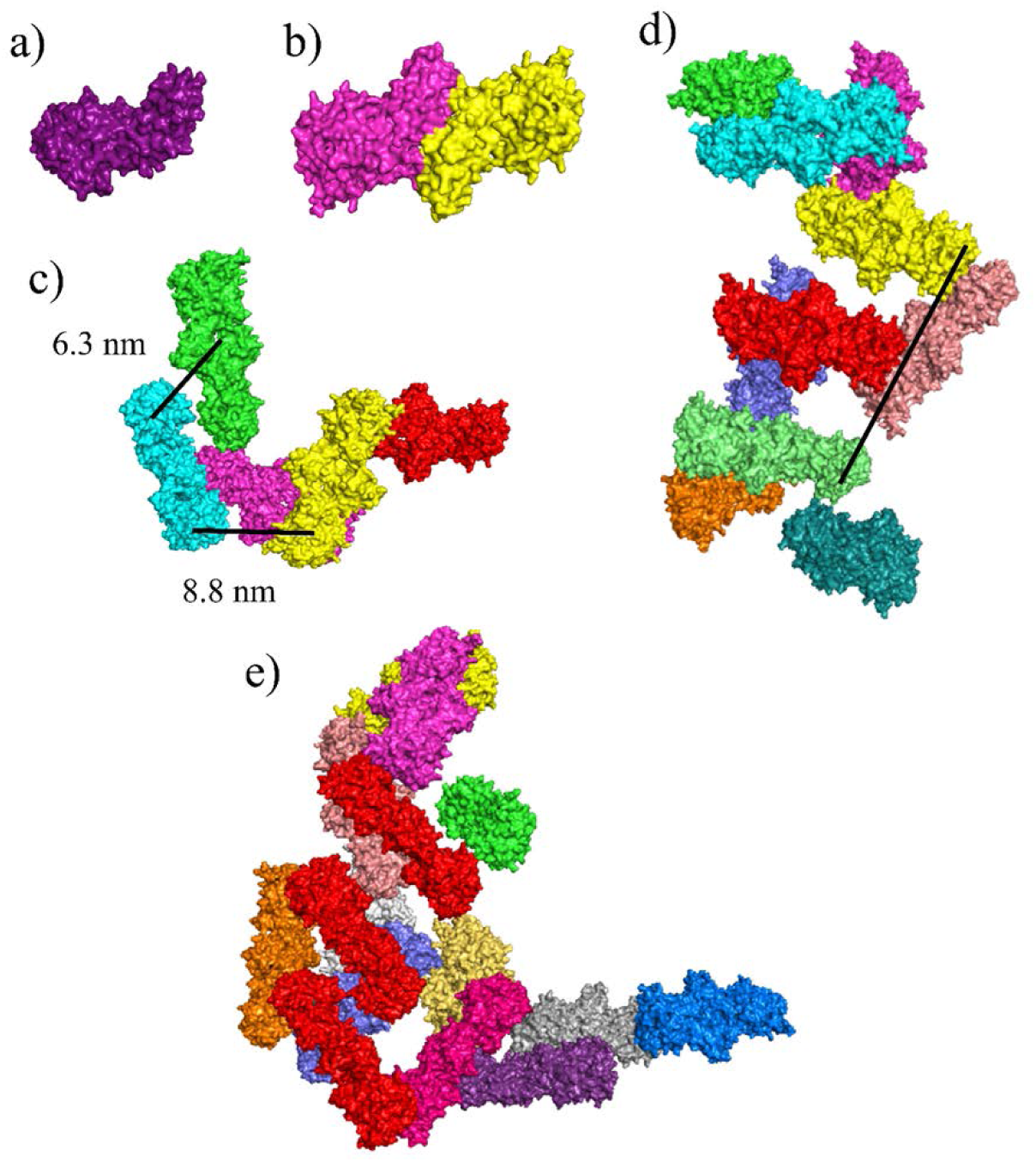
Structural models for the different states of LplA and the gel formation process as determined by SAXS. **a**. LplA monomer in solution. **b**. Dimer formed with H267 contacts between two monomer units. **c**. Pre-gel state at 20°C with K143 contacts between five dimer building blocks. **d**. Final gel state for *c* = 20 mg/mL solution made of 10 dimers. **e**. Final gel state for *c* = 40 mg/mL solution made of 15 dimers. In the fully developed gel, dimers form an arrangement of interlinked ladders (black line in **d**; marked in red in **e**), in which parallel dimers act as ladder steps that are connected by another sub-set of dimers. The characteristic distance of 8.8 nm is the spacing between these ladder steps, while 6.3 nm is the nearest-neighbor distance between two inclined dimer units.

SAXS data of different mutants of LplA show that mutations in the C-terminal (R277A and H267A) inhibit or at least shift both the dimer formation process and the gel formation to higher temperatures (**Fig. 7d**; **Tab. S2**). This observation suggests that a) the C-terminal domain is essential for dimerization and b) dimerization is a necessary step in the gel formation process. Therefore, as shown in **Fig. 9b**, a dimer model is created using two *SREFLEX*-based monomers (**Fig. 9a**) with close contact between the residues H267 in the C-terminal of two monomer subunits, which describes the SAXS curve (**Fig. 8a**). The so-created dimers served as building blocks to describe the scattering profiles at the different steps of the gelation process (**Fig. 8a**). For modeling the SAXS data, direct contacts between the K143 residues of two dimers were employed, which are at the two end sides of the dimer, since the LplA mutant of this residue (K143A) is found to hamper gel formation within the given temperature range (**Fig. 7d**; **Tab. S2**). With this approach the characteristic correlation peaks can be well described. As suggested by the increasing forward scattering intensity, the number of dimer subunits forming the average ordered cluster of the gel structure increases with temperature and concentration. Using these specific numbers of dimers as building blocks, models for the pre-gel state and gel state under the different conditions in the phase diagram are created by fitting the SAXS curves individually using the program *SASREF*.

The resulting models of the pre-gel and gel states are shown in **Fig. 9c-e**. Already at *T* = 20°C, i.e. at the onset of gel formation, the model reveals an ordered arrangement of the dimeric subunits. For the fully evolved gel, the presented models suggest a structure of interconnected ladders, in which parallel dimers serve as the steps of the ladders, while the ladder posts are formed by the chains of dimers. This long-range assembly therefore exhibits a lamellar order present in the SAXS data. While these clusters of dimers serve as an effective average model to describe the solution structure, the model covers all essential aspects of the gel structure. Specifically, the presented arrangement allows one to understand the characteristic length scales of the *d*-spacing as the next-neighbor distance between two connected dimers (*d* = 6.3 nm) as well as the spacing between two steps of the ladder-like assembly (*d* =8.8 nm). Moreover, it can explain the presence of cavities observed in the TEM images.

**Fig. 7d** displays the SAXS curves collected on solutions of five LplA mutants at *T* = 12°C (within the monomer state) and *T* = 30°C (at which the wild-type protein already has formed the full gel network) at c = 40 mg mL^-1^. For the low temperature regime, all mutants exhibit the SAXS profile similar to that of the wild-type, which is also reflected in the similar overall parameters (**Tab. S1**), revealing that the mutations do not affect the overall shape of the protein. The larger size and molecular weights of K143A and P167A reflect the presence of a higher amount of diffuse aggregates present from the beginning in solution.

Upon temperature rise, clear differences appear between the mutants. Mutant R277A is nearly unaffected, revealing that the mutant R277A in the C-terminal of LplA does inhibit dimer and thus gel formation. A similar result is observed for H267A, the other variant with mutation in this region for which only a weak formation of larger species is present. For the other mutants (K143A, R70A and P167A), with conserved C-terminal, the monomer-dimer transition is not inhibited but slightly shifted to higher temperatures. The size of diffuse aggregates / unordered clusters changes with the type of mutations (from 2 to 3 dimer unit equivalents), however, no ordered gel phase is formed. All these mutations in the tail part of LplA and thus of the dimers (K143A, R70A, R167A) clearly prevent the gel formation.

## Discussion

This study reports the ability of a natural non-structural protein, *E. coli* lipoate-protein ligase A (LplA), to self-assemble into a reversible hydrogel with a LCST behavior. The rheological properties, structure and formation mechanism of this unusual protein hydrogel are characterized. It is shown that the LCST phase transition and the gel properties can be modulated by pH, ions (buffers) and LplA concentration. The temperature and pH response ranges are close to physiological conditions, i.e. between 20-39°C, pH 5.5 −7.5. The stimuli-response ranges and LCST behavior are further varied by targeted mutations. Studies of the mutated LplA variants reveal also the key residues and domains, and thus the molecular mechanism(s) responsible for the gel formation, providing thus a solid basis for further engineering of LplA-based hydrogel for applications.

This study also provides a new model for the study of protein phase separation. The reversibility of the hydrogel indicates that LplA molecules are assembled in order, entailing a specific protein conformation and intermolecular interaction sites. Mutation studies have confirmed the essential roles of hydrogen bonding between the C-terminal domains and electrostatic interactions in the N-terminal domains for protein self-assembly. With the help of small-angle X-ray scattering and electron microscopy, we have obtained a more detailed insight into the molecular assembly process of LplA. The protein undergoes the formation of dimers, pre-clusters, and final stable aggregates with characteristic sizes. On the sub-molecular scale, we also observed the primary forms of protein aggregation, protein droplets, and synaptic-shaped gel networks. Since protein hydrogels have tunable properties, understanding the self-assembly mechanism of LplA will help to develop functional protein hydrogels with better performance in the future.

Last but not least, LplA is the first enzyme capable of forming hydrogel in a non-denatured state without the aid of crosslinking agents, while previously reported hydrogels made of globular proteins, such as bovine serum albumin (Khanna et al. 2021), ovalbumin and YajC-CT (Fang et al. 2011), are formed by irreversible protein amyloidization. *E. coli* LplA has been shown to be of great importance for C1 and energy metabolisms *in vivo*. It has an indispensable role in posttranslational modification of H-protein of the glycine cleavage system and acyltransferase subunit (E2) of the α-keto-acid dehydrogenase multi-enzyme complexes (Cronan 2016). Our preliminary study showed encouragingly that LplA remains its catalytic function to H-protein lipoylation of the glycine cleavage system at gel state. Thus, the unique catalytic activity of LplA hydrogel makes it an outstanding candidate as tissue regeneration material and biosensor *in vivo*. In addition, *E. coli* LplA is also a robust enzyme that has broad substrate specificity and has become a very useful tool for applications using *in vivo* protein labeling with fluorophores (Green et al. 1995; Liu et al. 2014). Recently, LplA has been engineered for site-specific antibody conjugates for sequential and orthogonal drug delivery and release (Thornlow et al. 2019). Thus, it would be of great interest to examine if LplA can be used to realize sol-gel transition inside cells or tissues. This would lead to paradigm changes in intracellular sensing, cell imaging and drug release. Furthermore, protein phase separation (Boeynaems et al. 2018) and engineering of intracellular protein condensate (Feng et al. 2019) are emerging as an exciting research area for understanding and controlling cellular compartments inside cells and for reengineering biological function in living cells. A realization of tunable LplA hydrogel inside living cells would allow engineering of intracellular condensates using a catalytically active protein instead of disordered proteins as presently dominating in the literature (Dzuricky et al. 2020).

## Methods

### Chemicals and Reagents

NaCl, Tris, and HCl were all of analytical grade and purchased from Sinopharm Chemical Reagent Co. LTD (Beijing, China). ATP, ADP, AMP and lipoic acid were purchased from Sigma-Aldrich (Shanghai, China). The bicinchoninic acid (BCA) Protein Assay Kit was purchased from Beijing Solarbio Science & Technology Co. LTD (Beijing, China). Chemically competent cells of *E. coli* TOP10 and *E. coli* BL21(DE3) were purchased from Weidishengwu Ltd. (Beijing, China). In-fusion cloning was used for the ligation of sequence fragments to vector with the In-fusion HD Cloning Kit (Clontech Laboratories, Inc, US). Site-directed mutations were accomplished according to the protocol provided in a Mut Express II Fast Mutagenesis kit V2 (Vazyme, Jiangsu, China). Luria-Bertani (LB) liquid medium (tryptone 10 g/L, yeast extract 5 g/L and NaCl 10 g/L) and solid medium (1.5% agar) with kanamycin (100 μg/mL), ampicillin (100 μg/mL) were used for transformation, screening, and cell growth.

### Protein Expression and Purification

The gene encoding *E. coli* LplA was constructed in pET-28a(+) vectors by In-fusion cloning. Site-directed mutations including R120A, K143A, K175A, S252A, H267A, D269A, R272A, R277A and Q279A were accomplished according to the protocol provided in a Mut Express II Fast Mutagenesis kit V2 (Vazyme, Jiangsu, China). Proteins were expressed in *E. coli* BL21(DE3). Recombinant cells were incubated at 37°C in Luria-Bertani medium containing 50 μg/mL kanamycin until the OD_600_ reached about 0.6, and 0.2 mM Isopropyl-β-D-Thiogalactoside (IPTG) was added to induce protein expression for 12 h at 30°C. Cells were then harvested by centrifugation at 10,000×g, 4°C for 10 min. Cell pellets were resuspended in buffer A (300 mM NaCl, 50 mM Tris-HCl, pH 7.5) and lysed by high-pressure homogenization. The lysed samples were centrifuged, and the supernatants were loaded onto and purified by a Ni^2+^ chelating Sepharose Fast Flow column (GE Healthcare), using an ÄKTA purifier (GE Healthcare). The recovered proteins were tested for purity on 12% SDS-PAGE.

### Hydrogel Preparation

Purified proteins were dialyzed against Tris-HCl buffer (50 mM, pH 7.5) using a 15 mL centrifugal filter unit with 30 kDa molecular weight cutoff (Amicon Ultra-15, Millipore) at 4°C and 2,250×g, after which the protein concentrations were assessed using a BCA Protein Quantitation Kit (Solarbio, Beijing, China). Samples of the purified proteins were diluted with the appropriate ice-cold buffer to 40 mg/mL, added into glass vials and placed at room temperature (20°C) for 5 minutes to observe the formation of hydrogels.

### Determination of enzyme activity for lipoylation

The reaction system for enzyme activity determination is given in **Table 1**. The enzyme activities of wild type LpLA and its mutant R277D were measured at a low concentration of 0.8 mg/mL (no gelation) and a relatively high concentration of 40 mg/mL (wild-type LplA in a gel state), respectively. The reaction was started with the addition of ATP, incubated at 30 °C for a certain time period, and then terminated by heating for 90s in boiling water to completely denature and precipitate LplA. The amount of lipoylated H-protein in the mixture was determined using HPLC as described in a previous study (Zhang et al. 2020b).

**Table 1.** The reaction system for enzyme activity determination

| Component | Solution state | Hydrogel state |
| --- | --- | --- |
| R-(+)-lipoic acid | 0.5 mM |  |
| ATP | 1.0 mM |  |
| MgCl<sub>2</sub> | 1.0 mM |  |
| DTT | 0.1 mM |  |
| H<sub>apo</sub> | 10 µM | 100 µM |
| Wild type LpLA or mutant R277D | 0.8 mg/mL (20 µM) | 40mg/mL (1.0 mM) |

### Dynamic Rheology Characterization

The rheological properties of LplA hydrogel were measured using a rotational rheometer instrument (AR2000ex, TA Instruments). LplA was dissolved in ice-cold Tris-HCl buffer at the concentration of 1.0 mM. The prepared LplA solutions were transferred to the sandblasted plate surfaces of the remoter previously set at 6°C and stabilized for 3 min. Anti-volatile rings were applied in order to prevent evaporative loss of solvent from the sample during the measurements. Strain sweeps and frequency sweeps were performed at 0.1% strain amplitude and 1.25 rad/s frequency, respectively. And temperature sweeps were performed at 1.25 rad/s frequency and 0.1% strain. In the continuous ramp of shear rate experiment, the shear rate is increased from 0.01 s^-1^ to 10^−1^ in 10 minutes, and continued with the performance of the reverse process. Temperature sweeps were also performed on the hydrogel formation of LplA mutants (R120A, K143A, K175A, S252A, H267A, D269A, R272A, R277A and Q279A)

### Electrostatic Potential Surface Reconstruction

Two crystal structures (PDB ID: 3A7R and 1×2H) of LplA were selected for the reconstruction of the electrostatic potential surface of LplA. The former (3A7R) was usually treated as a stretched conformation state of LplA, while the latter (1×2H) was considered as a bending conformation. The electrostatic potential surface of the protein was calculated and visualized by the plug-in APBS Tools (Baker et al. 2001) of PyMOL 2.1.0 (DeLano 2002). At first, the monomeric protein structures were extracted from the PDB files, with water and ligand removed in advance. Then PDB2PQR web server (Dolinsky et al. 2004) was employed to convert PDB-format structural information into PQR-format parameterized files, which were inputs for continuum solvation calculations in APBS. Finally, potential isosurfaces were obtained using the following parameters; 150 mM ionic strength, solute dielectric of 2.0, and solvent dielectric of 78.0.

### Electron Microscopy

Transmission electron microscopy (TEM) was performed on a Hitachi HT7700 TEM at 100.0 kV accelerating voltage and magnifications ranging from 30,000× to 50,000×. LpLA samples at a concentration of 40 mg/mL were negatively stained with 1% phosphotungstic acid in deionized H2O on carbon-coated 400 mesh Cu grids. The specimens were dried overnight before imaging.

Scanning electron microscopy (SEM) was performed on a HITACHI S-4700 SEM operating at 20 kV. Ice-cold hydrogel sample (*c* = 40 mg/mL) were left at room temperature for 10 min to ensure that the gel network is fully formed. Then, the samples were subjected to shock-freezing by liquid nitrogen, followed by lyophilization. After drying, the sample was carefully transferred onto sample stage covered with carbon adhesive, and sputtered with gold for 60 s.

### Small Angle X-ray Scattering (SAXS) Measurements

Small angle X-ray scattering (SAXS) measurements on the wild type LplA and five of its mutants (R277A, H267A, K143A, R70A and P167A) were performed in two separate experimental sessions at the BioSAXS beamline P12, EMBL/DESY, Hamburg, Germany (Blanchet et al. 2015) using the 150 µm (v) x 250 µm (h) beam at an X-ray energy of E = 10 keV (wavelength λ = 0.124 nm).

Sample solutions were automatically loaded into an in vacuum quartz capillary (inner diameter: 1.7 mm (first session); 0.9 mm (second session) (Schroer et al. 2018)) using the robotic P12 sample changer (Round et al. 2015) with continuous sample flow during data collection. Before and after the protein solutions, the corresponding buffer solutions were measured.

The measurements were performed over a temperature range from 8°C to 40°C with steps of 2°C and 5°C. For each temperature point, sample and buffer were freshly loaded to avoid beam-induced radiation damage. Two-dimensional SAXS patterns were recorded using a PILATUS 6M pixel detector at a sample-detector-distance of 3 m, covering the range of momentum transfer *s* = 0.04 – 7.0 nm-1 (*s* = 4π/λ sin(Θ), where 2Θ is the scattering angle). SAXS data were recorded as a sequential set of 20 images for every 50 ms (1st session) and 40 images every 100 ms (2nd session) of exposure. The experimental parameters for the different samples are summarized in **SI Tab. 1**.

The recorded SAXS patterns were azimuthally averaged (masking inter-module segments of the detector as well as the shadows of the beam stop and the flight tube), normalized to the transmitted beam, checked for the absence of radiation damage, averaged accordingly, and the scattering signal from the corresponding buffers was subtracted from the 1D-SAXS profiles of the proteins, all done by the P12 beamline SASFLOW pipeline (Franke et al. 2012). The resulting difference curves were further analyzed using the ATSAS software package (Franke et al. 2017).

From the SAXS profiles, the overall parameters (radius of gyration Rg, molecular weight MW) of the proteins were determined following the standard procedures (Svergun et al. 2013; Hajizadeh et al. 2018). To determine if the solution structure can be described by the high-resolution models of LplA, scattering profiles from these models (PDB ID: 3A7R and 1×2H) were computed by CRYSOL (Svergun et al. 1995). The monomer model fitting the SAXS data was generated using SREFLEX (Panjkovich and Svergun 2016).

To analyze the structure of aggregates prior to gel formation as well as of the gel network, rigid body modeling of the aggregates and ordered gel using monomers and dimers as building blocks was performed using SASREF (Petoukhov and Svergun 2005). Based on the results of the measurements, specific contacts between the building blocks were used as boundary conditions for the SASREF analysis.

## Support Information

**Table S1.**
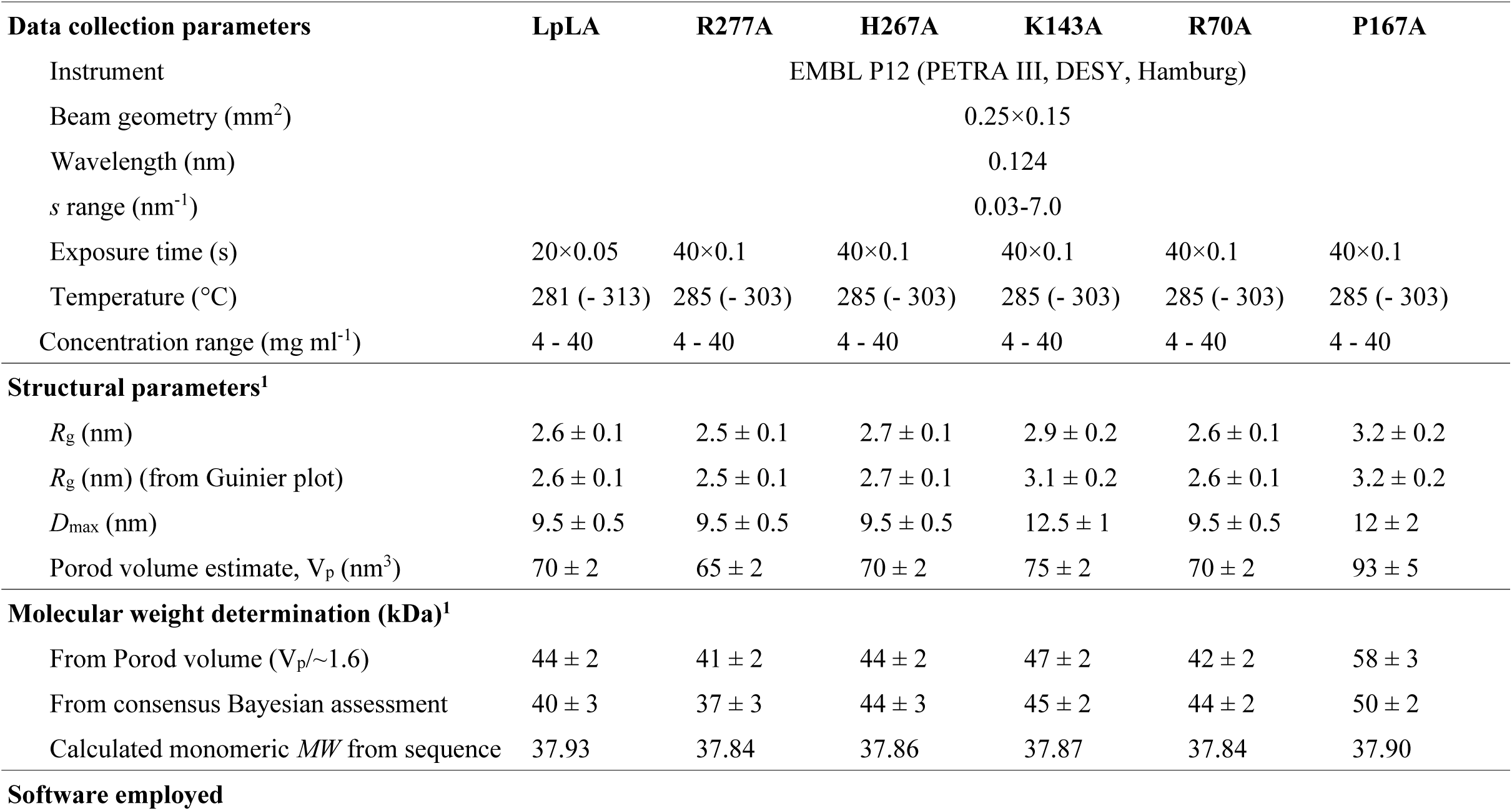
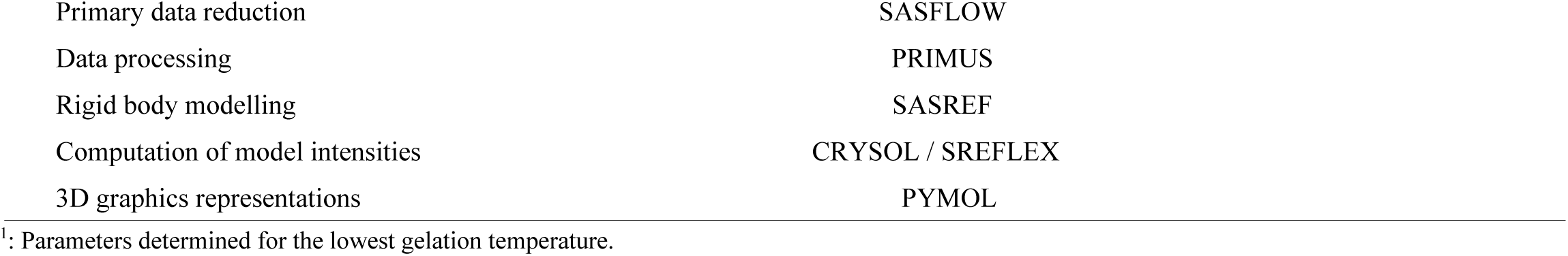
Data collection and structural parameters.

| **Data collection parameters** | **LpLA** | **R277A** | **H267A** | **K143A** | **R70A** | **P167A** |
| --- | --- | --- | --- | --- | --- | --- |
| Instrument | EMBL P12 (PETRA III, DESY, Hamburg) |  |  |  |  |  |
| Beam geometry (mm<sup>2</sup>) | 0.25×0.15 |  |  |  |  |  |
| Wavelength (nm) | 0.124 |  |  |  |  |  |
| *s* range (nm<sup>-1</sup>) | 0.03-7.0 |  |  |  |  |  |
| Exposure time (s) | 20×0.05 | 40×0.1 | 40×0.1 | 40×0.1 | 40×0.1 | 40×0.1 |
| Temperature (°C) | 281 (- 313) | 285 (- 303) | 285 (- 303) | 285 (- 303) | 285 (- 303) | 285 (- 303) |
| Concentration range (mg ml<sup>-1</sup>) | 4 - 40 | 4 - 40 | 4 - 40 | 4 - 40 | 4 - 40 | 4 - 40 |
| **Structural parameters<sup>1</sup>** |  |  |  |  |  |  |
| *R<sub>g</sub>* (nm) | 2.6 ± 0.1 | 2.5 ± 0.1 | 2.7 ± 0.1 | 2.9 ± 0.2 | 2.6 ± 0.1 | 3.2 ± 0.2 |
| *R<sub>g</sub>* (nm) (from Guinier plot) | 2.6 ± 0.1 | 2.5 ± 0.1 | 2.7 ± 0.1 | 3.1 ± 0.2 | 2.6 ± 0.1 | 3.2 ± 0.2 |
| *D<sub>max</sub>* (nm) | 9.5 ± 0.5 | 9.5 ± 0.5 | 9.5 ± 0.5 | 12.5 ± 1 | 9.5 ± 0.5 | 12 ± 2 |
| Porod volume estimate, *V<sub>p</sub>* (nm<sup>3</sup>) | 70 ± 2 | 65 ± 2 | 70 ± 2 | 75 ± 2 | 70 ± 2 | 93 ± 5 |
| **Molecular weight determination (kDa)<sup>1</sup>** |  |  |  |  |  |  |
| From Porod volume (*V<sub>p</sub>*/~1.6) | 44 ± 2 | 41 ± 2 | 44 ± 2 | 47 ± 2 | 42 ± 2 | 58 ± 3 |
| From consensus Bayesian assessment | 40 ± 3 | 37 ± 3 | 44 ± 3 | 45 ± 2 | 44 ± 2 | 50 ± 2 |
| Calculated monomeric *MW* from sequence | 37.93 | 37.84 | 37.86 | 37.87 | 37.84 | 37.90 |
| **Software employed** |  |  |  |  |  |  |

|  |  |
| --- | --- |
| Primary data reduction | SASFLOW |
| Data processing | PRIMUS |
| Rigid body modelling | SASREF |
| Computation of model intensities | CRY SOL / SREFLEX |
| 3D graphics representations | PYMOL |

<sup>1</sup>: Parameters determined for the lowest gelation temperature.

**Table S2.**
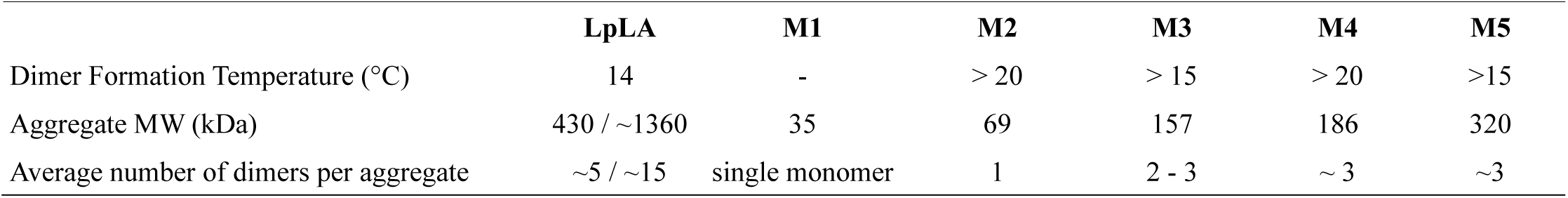
Dimer formation temperature, molecular weight & average number of dimers within the aggregates / gel cluster (*c* = 40 mg/mL). For wildtype, at 16°C (before onset of gel formation) and 35°C (full gel); for mutants at 30°C.

